# Warning SINEs: *Alu* elements, evolution of the human brain, and the spectrum of neurological disease

**DOI:** 10.1101/230367

**Authors:** Peter A. Larsen, Kelsie E. Hunnicutt, Roxanne J. Larsen, Anne D. Yoder, Ann M. Saunders

## Abstract

*Alu* elements are a highly successful family of primate-specific retrotransposons that have fundamentally shaped primate evolution, including the evolution of our own species. *Alu*s play critical roles in the formation of neurological networks and the epigenetic regulation of biochemical processes throughout the central nervous system (CNS), and thus are hypothesized to have contributed to the origin of human cognition. Despite the benefits that *Alus* provide, deleterious *Alu* activity is associated with a number of neurological and neurodegenerative disorders. In particular, neurological networks are potentially vulnerable to the epigenetic dysregulation of *Alu* elements operating across the suite of nuclear-encoded mitochondrial genes that are critical for both mitochondrial and CNS function. Here, we highlight the beneficial neurological aspects of *Alu* elements as well as their potential to cause disease by disrupting key cellular processes across the CNS. We identify at least 37 neurological and neurodegenerative disorders wherein deleterious *Alu* activity has been implicated as a contributing factor for the manifestation of disease and, for many of these disorders, this activity is operating on genes that are essential for proper mitochondrial function. We conclude that the epigenetic dysregulation of *Alu* elements can ultimately disrupt mitochondrial homeostasis within the CNS. This mechanism is a plausible source for the incipient neuronal stress that is consistently observed across a spectrum of sporadic neurological and neurodegenerative disorders.

**List of Abbreviations:**

A-to-I: adenosine-to-inosine
AD: Alzheimer’s Disease
ADAR: adenosine deaminase acting on RNA
ALS: Amyotrophic Lateral Sclerosis
AMPA: α-amino-3-hydroxy-5methyl-4-isoxazole propionate
APP: amyloid precursor protein
circRNAs: circular RNAs
CNS: central nervous system
FLAM: free left *Alu* monomer
LINE: long interspersed element
L1: long interspersed element-1
LTR: long-terminal repeat
mRNA: messenger RNA
PD: Parkinson’s Disease
pre-mRNA: precursor messenger RNA
SEDs: super-enhancer domains
SINE: short-interspersed element
TADs: topologically associating domains
TOMM: translocase of outer mitochondrial membrane

## Introduction

Retrotransposons are mobile genetic elements that utilize an RNA intermediate to copy and paste themselves throughout the genome. There are two primary groups of retrotransposons, those having long-terminal repeats (LTRs) and those without (non-LTR) (Cordaux and Batzer 2009). In the human genome, non-LTR retrotransposons consist of long interspersed elements (LINEs) and short interspersed elements (SINEs) and these collectively account for a remarkable ~33% of total genome sequence (Cordaux and Batzer 2009). *Alu* elements are primate-specific SINEs that are approximately 300 nucleotides in length and are abundant in the human genome, with over 1.3 million elements accounting for at least 11% of overall DNA sequence (Deininger et al. 2003; Hancks and Kazazian 2016). Although once considered to be useless ‘junk DNA’, the prevalence, diversity, and non-random distribution of *Alu* elements across primate genomes is suggestive of a functional advantage. Indeed, a large body of evidence documents that *Alu* elements have directly influenced primate evolution by facilitating genome innovation through: novel gene formation, elevated transcriptional diversity, long non-coding RNA and microRNA evolution (including circular RNAs), transcriptional regulation, and creation of novel response elements (Vansant and Reynolds 1995; Norris et al. 1995; Britten 1997; Lev-Maor et al. 2003; Polak and Domany 2006; Laperriere et al. 2007; Lin et al. 2008, 2016; Lehnert et al. 2009; Cordaux and Batzer 2009; Shen et al. 2011; Jeck et al. 2013; Töhönen et al. 2015; Luco 2016; Chen and Yang 2017). Moreover, *Alus* fundamentally alter the three-dimensional architecture and spatial organization of primate genomes by defining the boundaries of chromatin interaction domains (i.e., topologically associating domains (TADs); Dixon et al. 2012). A growing body of evidence indicates that genome architecture has a direct influence on biological function, and the observation that *Alus* are enriched within both TADs and super-enhancer domains (SEDs) supports the hypothesis that *Alus* directly influence a wide range of critically important processes within primates across multiple levels, from overall genome stability to tissue-specific gene regulation (Huda et al. 2009; Dixon et al. 2012; Soibam 2017; Glinsky 2018). In light of the functional benefits that *Alus* provide primates, it is interesting to note that *Alu* retrotransposition events occurred at an estimated 15-fold higher rate in the human, chimpanzee, and bonobo lineage (as compared to other great apes) and a 2.2-fold higher rate in humans when compared to chimpanzee and bonobo (Hedges et al. 2004; Prüfer et al. 2012; Hormozdiari et al. 2013). These evolutionary patterns indicate that positive selection is acting to maintain *Alu* elements in primate genomes, especially within humans (Mattick and Mehler 2008; Tsirigos and Rigoutsos 2009).

One of the most fascinating and biologically important aspects of *Alu* elements is that they serve an important role in the formation and function of the brain connectome (Oliver and Greene 2011; Li and Church 2013; Smalheiser 2014; Sakurai et al. 2014; Prendergast et al. 2014; Linker et al. 2017; Bitar and Barry 2018). Many lines of evidence connect *Alu* elements with neurogenesis and critical neuronal biochemical processes, including: somatic retrotransposition in developing neurons (in parallel to L1 retrotransposition; Baillie et al. 2011; Kurnosov et al. 2015), formation of regulatory circRNAs that are enriched in the central nervous system (CNS) and concentrated at synapses (Jeck et al. 2013; Rybak-Wolf et al. 2015; Chen and Schuman 2016; Floris et al. 2017), regulation of genes that are essential for proper neuron function (e.g. *ACE*, *SMN1*, *SMN2*, *SLC6A4*; Wu et al. 2013; Ottesen et al. 2017; Schneider et al. 2017), and elevated adenosine-to-inosine (A-to-I) RNA editing in the brain (Mehler and Mattick 2007; Kurnosov et al. 2015; Behm and Öhman 2016). In particular, epigenetic A-to-I editing plays a significant role in mediating neuronal gene expression pathways (Tariq and Jantsch 2012) with *Alus* serving as the primary target for RNA editing in primates (Picardi et al. 2015; Behm and Öhman 2016). Beyond RNA editing mechanisms, human neuronal gene pathways are regulated by noncoding RNAs originating from *Alu* elements (e.g., BC200 and NDM29) and specific *Alu* subfamilies contain retinoic acid response elements which help to regulate neural patterning, differentiation, and axon outgrowth (Vansant and Reynolds 1995; Laperriere et al. 2007; Maden 2007; Castelnuovo et al. 2010; Smalheiser 2014). Moreover, recent discoveries indicate *Alu* elements underlie the formation of a vast number of human-specific circRNAs that are hypothesized to play important roles in neurological gene expression pathways (Jeck et al. 2013; Rybak-Wolf et al. 2015; Chen and Schuman 2016; Dong et al. 2017). There is a deep connection between *Alus* and the formation and function of primate neurological networks, and this has led to the hypothesis that *Alu* elements were essential for development of the transcriptional diversity and regulation required for the genesis of human cognitive function (Mattick and Mehler 2008; Oliver and Greene 2011; Li and Church 2013; Sakurai et al. 2014).

Despite the functional benefits that *Alus* have provided primate genomes, *Alu* elements can disrupt gene expression and function through many pathways (Figure 1; Deininger and Batzer 1999; Deininger 2011; Tarallo et al. 2012; Ade et al. 2013; Elbarbary et al. 2016; Varizhuk et al. 2016). For this reason, the genome tightly regulates *Alus* using both DNA methylation and histone (H3K9 methylation) modification in order to control their expression and *de novo* retrotransposition (Varshney et al. 2015; Elbarbary et al. 2016; Mita and Boeke 2016) and there is mounting evidence indicating that the loss of these epigenetic control mechanisms (due to aging, cellular senescence, environmental factors and stress) contributes to many forms of cancer, diabetes, osteoporosis, and several mental and neurodegenerative disorders (Szpakowski et al. 2009; Belancio et al. 2010; Muotri et al. 2010; Jintaridth et al. 2013; Dannlowski et al. 2014; Erwin et al. 2014; Bundo et al. 2014; Sun et al. 2014; Goodier 2016; Neven et al. 2016; Bedrosian et al. 2016; Shpyleva et al. 2017; Thongsroy et al. 2017). With respect to deleterious *Alu* pathways and neurological disease, there are at least 37 mental and neurodegenerative disorders wherein *Alu* elements are hypothesized to disrupt key cellular processes, thereby resulting in or contributing to the diseased state (Table 1).

**Figure 1.**
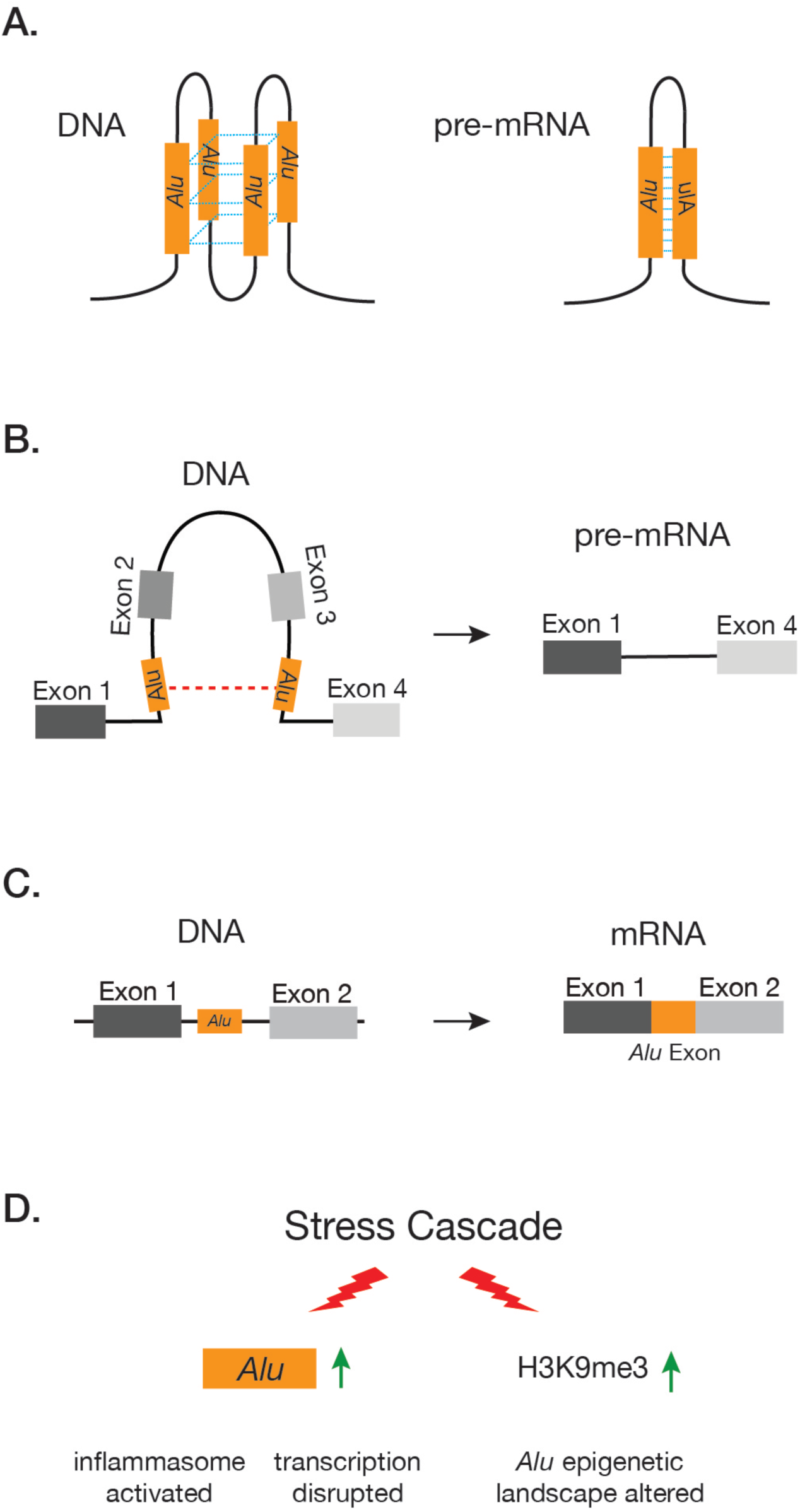
Select mechanisms whereby *Alu* elements can alter gene expression and function (also see Elbarbary et al. 2016). **A:** Sequence homology and orientation of *Alu* elements contributes to the formation of distinct secondary structures in both DNA and RNA. DNA *Alu* G-quadruplex structures can alter transcription kinetics (Varizhuk et al. 2016) and pre-mRNA *Alu* binding forms stem-loop structures that are the primary site for A-to-I editing (see Figure 2). **B:** Recombination of intra-gene *Alu* elements resulting in exon deletion. **C:** Exonification of intronic *Alus* contributing to the production of alternative mRNAs. **D:** Environmental or traumatic stress cascades resulting in increased expression of *Alu* RNAs that contribute to inflammation (Li and Schmid 2001; Tarallo et al. 2012; Hunter et al. 2015; Lapp et al. 2016), the disruption of global gene transcription through Pol II binding (Mariner et al. 2008), and an increase of H3K9 histone methylation that alters *Alu* epigenetic pathways (Varshney et al. 2015; Lapp and Hunter 2016; Larsen et al. 2017).

**Table 1.** Genes associated with neurological and neurodegenerative disorders wherein deleterious *Alu* activity has been documented experimentally or is hypothesized to disrupt gene function. Gene names in bold identify genes essential for mitochondrial function and transport and/or are associated with mitochondrial abnormalities (sensu Dawson et al. 1995, Calvo et al. 2015, Zempel and Mandelkow 2015, Bhattachargee et al. 2016, Chong-Chong et al. 2016, Checler et al. 2017, Johnson et al. 2017, Wang et al. 2013). For additional *Alu* associated diseases see Hancks and Kazazian (2016) and Payer et al. (2017). Asterisks identify genes where mutations result in dysregulation of *Alu* elements.

| Gene Name | Disorder | *Alu* Mechanism of Disruption | Reference |
| --- | --- | --- | --- |
| **ABCD1** | Adrenoleukodystrophy | Deletion events | Kutsche et al. 2002 |
| **ACAT1 (T2)** | Mitochondrial acetoacetyl-CoA thiolase deficiency | Deletion event | Zhang et al. 2006 |
| ACE | Alzheimer's disease | Insertion events | Wu et al. 2013 |
| ADAR2 | Glioma | Exonization | Li et al. 2015 |
| **ALDH7A1** | Pyridoxine-dependent epilepsy | Recombination | Mefford et al. 2015 |
| ALMS1 | Alström syndrome | Insertion event | Taşkesen et al. 2012 |
| APOB | Hypobetalipoproteinemia | Recombination | Huang et al. 1989 |
| **ATP5J** | Alzheimer's disease | Duplication | Antonell et al. 2012 |
| **ATP7A** | Menkes disease | Insertion event | Gu et al. 2007; Bhattacharjee et al. 2016 |
| **ATP7B** | Wilson's disease | Alternative splicing | Mameli et al. 2015 |
| **C9orf72** | ALS, FTLD | Loss of epigenetic control, elevated *Alu* transcripts | Prudencio et al. 2017 |
| CHD7 | CHARGE syndrome | Deletion | Udaka et al. 2007 |
| **CLN3** | Batten disease | Deletion | Lerner et al. 1995 |
| COL4A5 | Alport syndrome | Deletion and exonization | Nozu et al. 2014 |
| DICER1* | Age-related macular degeneration | *Alu* RNA build-up with reduced DICER1 activity | Kaneko et al. 2011; Kim et al. 2014 |
| **FXN** | Friedreich's Ataxia | *Alu* repeat expansion, alternative splicing events | Pandolfo 2006 |
| **GK** | Glycerol kinase deficiency | Insertion event | Zhang et al. 2000 |
| GLA | Fabry disease | Deletion event | Dobrovolny et al. 2011 |
| HPRT | Lesch-Nyhan disease | Recombination | Brooks et al. 2001 |
| **HMBS** | Acute intermittent prophyria | Insertion event | Mustajoki et al. 1999 |
| LPL | Lipoprotein lipase deficiency | Complex deletion-insertion | Okubo et al. 2007 |
| **MFN2 (CMT2a)** | Charcot-Marie-Tooth type 2A | Copy number variants | Pehlivan et al. 2016 |
| MPO | Alzheimer's disease | *Alu* hormone response variant; estrogen dysregulation | Reynolds et al. 1999 |
| **NDUFS2** | Leigh Syndrome | Exonization | Larsen et al. 2017 |
| NF1 | Neurofibromatosis type I | Deletion and chimeric gene fusion | Wimmer et al. 2011; Ferrari et al. 2017 |
| NFIX | Marshall-Smith syndrome | Deletions | Schanze et al. 2014 |
| **OPA1** | Autosomal Dominant Optic Atrophy | Alternative splicing events | Gallus et al. 2010 |
| **PARK2** | Parkinson's disease | Recombination | Morais et al. 2016 |
| **PARK7 (DJ-1)** | Parkinson's disease | Deletion | Bonifati et al. 2002 |
| **PDHA1** | Pyruvate Dehydrogenase Deficiency | Exonization | Larsen et al. 2017 |
| PIGL | CHIME syndrome | Deletion | Knight Johnson et al. 2017 |
| PMM2 | Congenital disorders of glycosylation type Ia | Complex deletion | Schollen et al. 2007 |
| POMT1 | Walker Warburg syndrome | Insertion | Bouchet et al. 2007 |
| **PSEN1** | Alzheimer's disease | Deletion | Le Guennec et al. 2017 |
| **PXMP2 (PMP22)** | Charcot-Marie-Tooth type 2A | *Alu-Alu*-mediated rearrangement | Choi et al. 2011; Gu et al. 2015 |
| **RP2 (NUDT19)** | X-linked retinitis pigmentosa | *Alu*-L1 recombination | Schwahn et al. 1998, Jiang et al. 2017 |
| **SLC6A4** | Depression, reduced hippocampal volume | Altered promoter methylation | Dannlowski et al. 2014 |
| **SLC25AC** | Intellectual disability | Deletion | Vandewalle et al. 2013 |
| **SLC30A6** | Alzheimer's disease, Dementia, ALS | Gene fusion event | Boone et al. 2014 |
| **SMN1** | Spinal muscular atrophy | Exonization, deletion events, circularization | Ottesen et al. 2017 |
| **SOD2** | Hyperglycemia | Repressed expression | Wang et al. 2016 |
| SOX10 | Waardenburg syndrome type 4 | Deletion | Bondurand et al. 2012 |
| **SPAST** | Autosomal-dominant spastic paraplegia 4 | Deletions, CNVs, gene fusion events | Boone et al. 2014 |
| **SPG7** | Hereditary spastic paraplegia | Deletion, recombination | Arnoldi et al. 2008; López et al. 2015 |
| SPG11 | Hereditary spastic paraplegia | Deletion | Conceição et al. 2012 |
| STAU1 | Myotonic Dystrophy Type 1 | Alternative splicing regulation | Bondy-Chorney et al. 2016 |
| **TDP-43** | ALS, frontotemporal lobar degeneration | Transposable element dysregulation | Li et al. 2012 |
| **TOMM40** | Late-Onset Alzheimer's Disease | *Alu* repeat expansion, putative alternative splicing events | Larsen et al. 2017 |
| TRIM37 | Mulibrey nanism | Deletion events | Jobic et al. 2017 |

Given the tight connection between *Alu* elements and the formation and function of the nervous system, it is likely that the dysregulation of *Alu* elements contributes to many sporadic or idiopathic neurological disorders observed across the global human population (Larsen et al. 2017). Here, we highlight both the beneficial neurological aspects of *Alu* elements as well as their potential to cause neurological disease. We focus on a novel hypothesis that identifies a potential epigenetic vulnerability to neurological networks that has likely escaped purifying selection. The *Alu* neurodegeneration hypothesis (sensu Larsen et al. 2017) posits that the epigenetic dysregulation of *Alu* elements ultimately serves to disrupt mitochondrial homeostasis in neurological networks, thereby setting the stage for increased neuronal stress and neurodegeneration. Given this hypothesis, it is noteworthy that many of the *Alu*-disrupted genes associated with neurological disorders are related to mitochondrial function and trafficking, including nuclear-encoded mitochondrial genes (i.e., mitonuclear) which help to regulate oxidative stress and metabolic processes in the CNS (Table 1). Mitochondrial dysfunction is implicated across the spectrum of neurological and neurodegenerative disorders that are observed in humans and this pattern is suggestive of a genetic vulnerability that has evolved in humans. Considering this, we begin by reviewing the integral role that *Alu* elements have played in human evolution through brain-specific epigenetic A-to-I RNA editing pathways and neurological network formation. Although these *Alu*-related processes are hypothesized to have contributed to the origin of human cognition, they are likely accompanied by age or stress-related epigenetic vulnerabilities to the CNS, with mitochondrial pathways being especially sensitive.

### *Alu* elements, A-to-I editing, and evolution of the human brain

*Alu* elements are non-randomly distributed throughout the genome. They occur most frequently within introns and are enriched within genes involved in metabolic, mitochondrial, cellular transport, and binding pathways (Grover et al. 2003; de Andrade et al. 2011; Larsen et al. 2017). *Alu* nucleotide sequences and lengths (~300 bp) are generally conserved (Batzer and Deininger 2002) and it is this seemingly simple aspect of *Alu* biology that is of monumental biological importance. When inserted within a gene at opposite orientations and at close proximity, *Alus* bind upon themselves post-transcriptionally, resulting in the formation of a duplex stem-loop structure that is stabilized by the *Alu* nucleotide sequence and length (Figure 2; Athanasiadis et al. 2004). These *Alu*-based secondary structures fundamentally alter the shape of pre-mRNA molecules and serve as the primary binding site for ADAR proteins, which bind to the double-stranded pre-mRNA duplex and edit adenosine (A) residues to inosine (I) thereby recoding pre-mRNAs (Figure 2). When operating in coding regions (either directly or indirectly), the translation machinery interprets the resulting I residues as guanosine (G) and this mechanism accounts, in part, for the incredible diversity observed in the human proteome that is not encoded within the original DNA sequence (Nishikura 2016). However, the vast majority of A-to-I editing operating on *Alu* elements occurs within pre-mRNA introns and 3’ UTRs and this can directly influence gene regulation and function in a surprising number of ways, including: the creation of novel splice donor and acceptor sites that result in *Alu* exonization and alternative gene splicing (Nishikura 2016); recoding of exons immediately adjacent to *Alus* (Daniel et al. 2014); disruption of RNAi pathways (Chen and Carmichael 2008); production of novel micro-RNA regulatory sites (Borchert et al. 2009); and increased nuclear retention of promiscuously edited mRNAs (Chen and Carmichael 2008).

**Figure 2.**
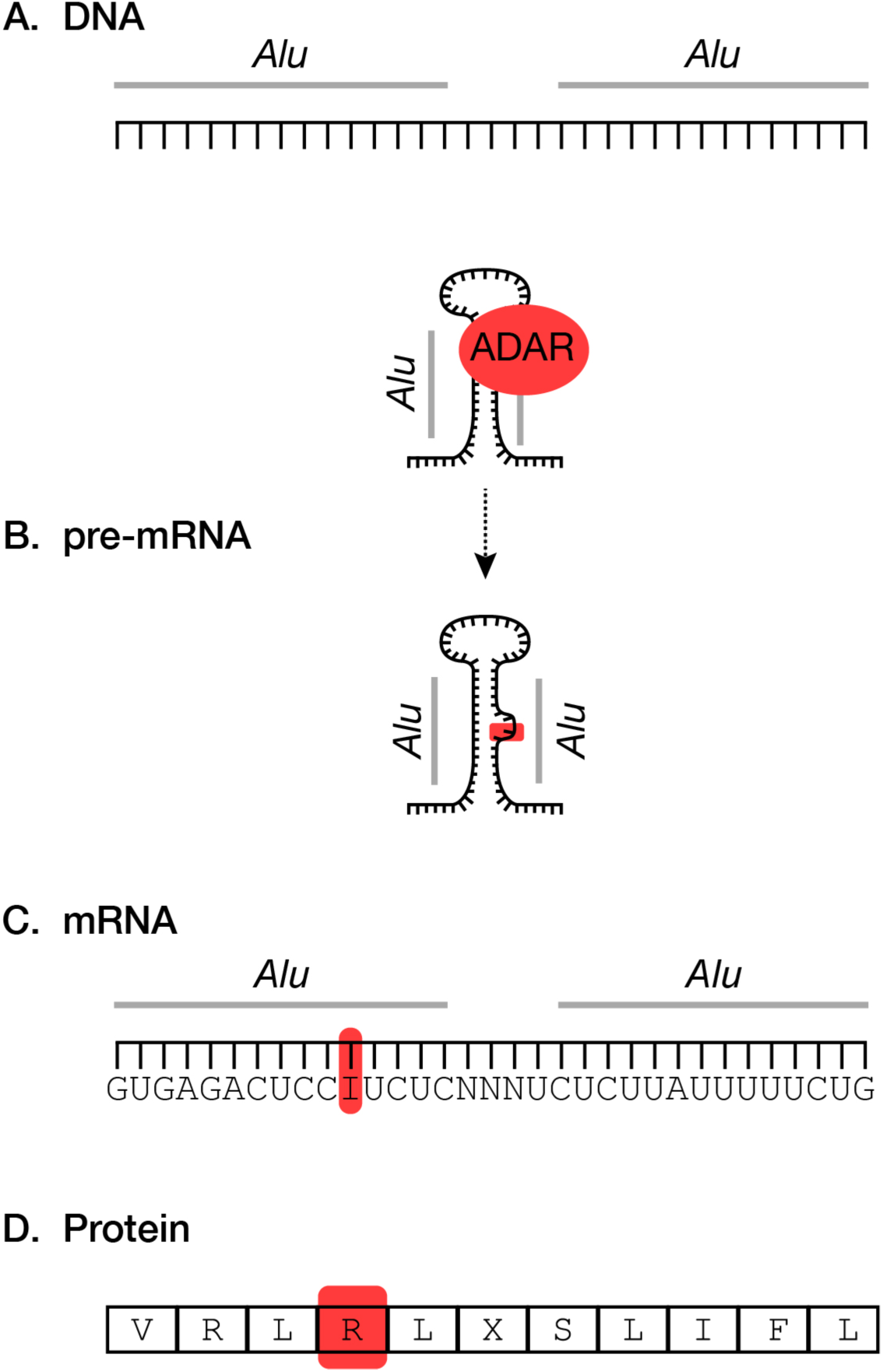
Intronic *Alu* elements located in close proximity (**A**) can bind to each other within pre-mRNAs (**B**) thereby producing a stable stem-loop secondary structure that is the primary substrate for A-to-I editing in primates. ADAR proteins bind to pre-mRNA *Alu* structures (**C**) and convert adenosine residues to inosine. If occurring in coding regions, the translation machinery then interprets the inosine residues as guanosine and this can contribute to amino acid changes and alternative protein conformations (**D**).

Although A-to-I editing plays an essential role in generating transcriptional diversity across eukaryotes, *Alu* elements provide primate-specific RNA editing opportunities. An example of this is found when comparing the rodent-specific SINE B1 family to *Alu*. Both B1 and *Alu* SINE families originated from 7SL RNA (Ullu and Tschudi 1984; Vassetzky et al. 2003) yet rodent-specific B1 elements are approximately half the length (~140 bp) of primate *Alus* (~300 bp) and have greater levels of intra-nucleotide variation. When considering the functional mechanics of A-to-I editing (Figure 2), the shorter lengths and more variable rodent B1 elements result in the formation of shorter and less-stable double-stranded stem-loop structures in rodent pre-mRNAs. Thus, the molecular properties that separate rodent B1 from primate *Alu* translate to key functional genomic differences that have influenced evolutionary processes within each lineage (Eisenberg et al. 2005; Neeman et al. 2006; Picardi et al. 2015; Tan et al. 2017).

A-to-I editing associated with *Alu* elements is perhaps one of the most functionally important yet underappreciated aspect of *Alu* biology. Approximately 90% of A-to-I editing within primate gene networks centers on *Alu* elements and this has fundamentally shaped primate evolution, including the evolution of our own species where A-to-I editing is estimated to occur at over 100 million sites in the human transcriptome (Bazak et al. 2014). Moreover, recent data supports a connection between A-to-I editing and neurological network formation, with elevated editing levels occurring throughout neurogenesis (Behm and Öhman 2016). Genes encoding for key neurological proteins involved in neurotransmission, neurogenesis, gliogenesis, and synaptogenesis are subject to enhanced A-to-I editing and thus a number of studies have hypothesized a strong link between A-to-I RNA editing pathways and brain development and function (Schmauss and Howe 2002; Mehler and Mattick 2007; Tan et al. 2009; Sakurai et al. 2014; Liscovitch et al. 2014; Behm and Öhman 2016; Hwang et al. 2016; Picardi et al. 2017b). A recent analysis of A-to-I editing in over 8,500 human samples identified tissue specific editing patterns with elevated editing levels in the brain, including unique patterns in the cerebellum (Tan et al. 2017). The vast majority of these neurologic A-to-I editing events are operating on *Alu* elements and, when combined with human-specific *Alu* evolution (Hedges et al. 2004; Cordaux and Batzer 2009; Prüfer et al. 2012; Hormozdiari et al. 2013), this observation serves as the foundation for the hypothesis that *Alu* elements and *Alu*-related pathways contributed to the evolution of enhanced human cognitive abilities (Mattick and Mehler 2008; Li and Church 2013).

Given the relationship between *Alu*-centric A-to-I editing and the formation and function of the CNS, it is important to expand upon the neuro-specific functions of ADAR proteins. Three ADAR proteins are identified (ADAR1, ADAR2, and ADAR3) and these proteins have distinct tissue-specific expression patterns (Picardi et al. 2015; Tan et al. 2017). ADAR1 and ADAR2 co-opt to regulate neuronal activity by editing key neurotransmitter receptors and ion channels in the CNS (Hood and Emeson 2011). Interestingly, both ADAR2 and ADAR3 have unique brain-specific expression patterns with ADAR2 being highly expressed in the brain and ADAR3 exclusively expressed in the brain (Mehler and Mattick 2007). Until recently the functional role of the brain-specific ADAR3 protein was largely unknown, however, Oakes et al. (2017) discovered that ADAR3 competes with ADAR2 to regulate glutamate receptor subunit B (*GRIA2*) A-to-I editing. The *GRIA2* protein forms a critical subunit of α-amino-3-hydroxy-5methyl-4-isoxazole propionate (AMPA) receptors, which regulate synaptic calcium and are involved with synaptic plasticity, memory, and learning (Wright and Vissel 2012). Remarkably, A-to-I editing of a specific adenosine nucleotide within *GRIA2* results in an amino acid change that alters the GluR-2 protein conformation, thus disrupting calcium permeability of the AMPA receptor and potentially contributing to epilepsy, amyotrophic lateral sclerosis (ALS), and schizophrenia (see Oakes et al. 2017). In light of Oakes et al. (2017), the brain-specific expression pattern of ADAR3 indicates that this protein helps to offset A-to-I editing by ADAR2, perhaps serving to mediate enhanced RNA editing processes throughout the CNS.

Considering the essential role that A-to-I editing processes play in the CNS, the dysregulation of these processes can have a profound impact on the stability of neurological networks (Mehler and Mattick 2007; Rice et al. 2012; Hwang et al. 2016). With respect to ADAR proteins, mutations within ADAR1 have been linked to Aicardi-Goutières syndrome (characterized by severe brain dysfunction; Rice et al. 2012) and *Alu*-related alternative splicing events of ADAR2 are linked to glioma (Li et al. 2015). Disruption of ADAR1 editing increases production of unedited RNAs which interact with MAV proteins in the outer mitochondrial membrane, ultimately serving to activate inflammatory response pathways (Bajad et al. 2017; Gallo et al. 2017) and perhaps providing a mechanism for inflammatory diseases of the CNS (Hofer and Campbell 2016). ADAR2 knockout mice display epileptic seizures and neuronal death caused by an influx of calcium owing to the disruption of *GRIA2* editing (see above). The interference of A-to-I editing processes associated with the *KCNA1* gene (encoding a protein essential for potassium regulation and neuron excitability) is hypothesized to underlie Episodic Ataxia Type-1 disorder, a disease of the CNS characterized by seizures, stress-induced ataxia, and myokymia (Ferrick-Kiddie et al. 2017). Moreover, a reduction of A-to-I editing has been observed within hippocampal tissues of Alzheimer’s brains versus healthy controls (Khermesh et al. 2016). From a broader perspective, the disruption of A-to-I editing processes across the CNS has been linked to a wide variety of mental and neurodegenerative disorders including major depression and suicide, epilepsy, schizophrenia, Alzheimer’s disease (AD), and ALS (Gurevich et al. 2002; Kawahara et al. 2004; Kwak and Kawahara 2005; Maas et al. 2006; Kubota-Sakashita et al. 2014; Khermesh et al. 2016; Weissmann et al. 2016; Gal-Mark et al. 2017).

### *Alu* elements, neurogenesis, and the human brain connectome

There is a strong connection between *Alu* A-to-I editing and the development and function of the brain, therefore it is impossible to disentangle *Alus* from the formation and function of neurologic networks (Mehler and Mattick 2007; Tan et al. 2009; Behm and Öhman 2016). It is estimated that the human brain is comprised of over 100 billion neurons that are organized into functional hubs or parcels collectively forming the brain connectome (Van Den Heuvel and Sporns 2013). Beyond major structures of the brain (e.g., cerebellum, frontal cortex, hippocampus, etc.), the existence of connectome parcels shared across unrelated individuals is indicative of an evolutionary conserved process underlying neurological network formation and operating throughout neurogenesis. For example, a recent study mapped the cerebral cortex using multi-modal magnetic resonance imaging and identified 180 connectome parcels that were largely shared across 210 healthy adults (Glasser et al. 2016). Understanding the molecular processes that contribute to the formation of the human brain connectome is essential for understanding the origin of human-specific neurological disorders and diseases observed across the global distribution of our species. This is especially true for neurodegenerative conditions that are hypothesized to originate in functional network hubs and progress along neuronal network connections (e.g., AD; Seeley 2017; Cope et al. 2018).

A growing body of evidence indicates that retrotransposons (including both LINEs and SINEs) are active throughout neurogenesis and contribute to mosaic neuron genomes that ultimately form the human brain connectome (Muotri et al. 2005; Erwin et al. 2014; Kurnosov et al. 2015; Evrony 2016; Paquola et al. 2016; Linker et al. 2017). Although somatic L1 retrotransposition events within developing neurons have received much attention, it is noteworthy that *Alu* retrotransposition occurs in parallel with L1 throughout neurogenesis (Baillie et al. 2011; Kurnosov et al. 2015), thus providing primate-specific aspects of neurologic network formation. Furthermore, there is evidence that unites *Alu* elements with retinoic acid regulation (Vansant and Reynolds 1995; Laperriere et al. 2007), which is essential for neuronal patterning and differentiation throughout neurogenesis and is a potential regulator of neuron regeneration (Maden 2007). Retinoic acid is vital for the establishment, maintenance, and repair of neuronal networks and, given the presence of retinoic acid response elements in *Alu* elements, it is possible that *Alu* activity during neurogenesis is connected to retinoic acid signaling processes.

Considering the *Alu* regulatory pathways discussed above, it is of great interest to note that retrotransposition of *Alu* elements is hypothesized to occur at elevated levels within the dentate gyrus of the hippocampus, the putative site of adult neurogenesis (Kurnosov et al. 2015). Moreover, A-to-I editing levels steadily increase as neural progenitor cells develop into adult neurons (Behm and Öhman 2016). These data indicate that at least two retrotransposon-centric processes (somatic retrotransposition of both LINES and SINES and enhanced A-to-I editing operating primarily on *Alu* elements) are major contributors to neurogenesis, perhaps serving to establish the neuronal and biochemical diversity that underlies the ~100 billion neuron brain connectome. Remarkably, emerging data suggests that a third *Alu*-centric process is associated with the formation and function of neurological networks, this being the production of circRNAs that are enriched in the brain and concentrated at synaptic junctions (Jeck et al. 2013; Rybak-Wolf et al. 2015; Chen and Schuman 2016). Identifying vulnerabilities to each of these retrotransposon-centric processes will likely contribute to the identification of novel mechanisms underlying mental disorders and neurologic disease and could lead to novel therapeutic interventions.

### Pathways to incipient neuronal stress and neurological disease

The disruption of *Alu*-centric epigenetic RNA editing processes is implicated across the entire spectrum of neurologic disorders (see above). In light of this observation, it is interesting to note that another, seemingly unrelated, feature of many neurological disorders is mitochondrial dysfunction (Lin and Beal 2006; Rugarli and Langer 2012; Gottschalk et al. 2014; Petschner et al. 2017). However, we have previously shown that mitonuclear genes are enriched with *Alu* elements when compared to random (Larsen et al. 2017), which is consistent with earlier observations regarding the non-random insertion of *Alu* elements into genes associated with transcriptionally active regions of the genome (Grover et al. 2003; de Andrade et al. 2011). Thus, it is likely that *Alu*-mediated gene regulatory processes are actively influencing mitonuclear gene expression, regulation, and protein function through the pathways discussed above and reviewed in Chen and Yang (2017). Knowing this, the dysregulation of epigenetic *Alu* regulatory pathways is a plausible source for mitochondrial stress and dysfunction, with the CNS being particularly vulnerable (Larsen et al. 2017). Such a mechanism could contribute to the initial activation of complex mitochondrial stress pathways and incipient neuronal stress associated with sporadic neurologic disorders (e.g., inflammation, immune response, mitophagy, etc.). Importantly, these processes would precede macroscopic pathologies such as protein aggregation and neuronal atrophy observed in neurodegenerative diseases (Swerdlow et al. 2010; Larsen et al. 2017; Swerdlow 2017).

The *Alu* neurodegeneration hypothesis (*sensu* Larsen et al. 2017) proposes a ‘double-edged sword’, whereby the beneficial *Alu*-related processes that underlie neuron diversity and function also have the potential to disrupt mitochondrial homeostasis across neurological networks through deleterious cascade events that are facilitated by eroding tissue-specific *Alu* epigenetic control mechanisms. The stability of the brain’s connectome and the entire CNS depends on healthy mitochondrial populations within neurons, astrocytes, microglia and supporting cells (Cai et al. 2011; Viader et al. 2011; Schwarz 2013; Jackson and Robinson 2017). Mitochondria play critical roles for a wide range of essential neuronal processes including glucose and lipid metabolism, metal ion biosynthesis, cellular trafficking along axons, neurotransmitter relay across synapses, and synaptic calcium homoeostasis (Schwarz 2013; Harbauer et al. 2014). Therefore, molecular mechanisms that are known to disrupt gene expression and protein folding of genes that are essential for mitochondrial function can ultimately disrupt neurological function.

Interference of mitochondrial dynamics across the CNS is consistently hypothesized to occur during the earliest stages of mental, neurological, and neurodegenerative disorders ranging from depression, epilepsy, and schizophrenia to ALS, AD, and Parkinson’s disease (PD; Lu 2009; Rezin et al. 2009; Kim et al. 2010; Coskun et al. 2012; Martin 2012; Gottschalk et al. 2014; Larsen et al. 2017; Flippo and Strack 2017; Petschner et al. 2017). Collectively, these disorders are estimated to impact approximately 250 million people globally, accounting for at least 10.2% of the global disease burden (GBD 2015 Neurologic Disorders Collaborator Group 2017). The occurrence of sporadic forms of human-specific neurologic disorders (e.g. non-familial schizophrenia, ALS, late-onset AD, PD, etc.) across the entire distribution of our species is suggestive of a common yet complex genetic mechanism that evolved in primates and is amplified in humans (Larsen et al. 2017). Considering this, we expand on the mitocentric view of idiopathic neurologic disease manifestation by reviewing the evidence that unites primate-specific *Alu* activity with incipient neurologic mitochondrial dysfunction.

Eukaryotic mitochondria are hypothesized to have originated from an endosymbiotic alphaproteobacterium which, over expansive evolutionary time, evolved in parallel with host genomes into the mitochondrial organelles that we observe today (Roger et al. 2017). The human mitochondrial genome encodes only 13 proteins yet it is estimated that human mitochondria depend on approximately ~2,000 genes encoded within the nuclear genome for their functionality (Calvo et al. 2015; Johnson et al. 2017). These mitonuclear genes are thus subject to deleterious *Alu* activity and *Alu*-related deleterious events have been linked to many neurologic and neurodegenerative disorders, including epilepsy, Wilson’s disease, Leigh syndrome, PD, ALS, and AD (Table 1 and references therein; Figure 3). When considering the incipient mitochondrial dysfunction observed across the spectrum of neurological neurodegenerative disorders, it is possible that tissue-specific epigenetic dysregulation of *Alu* elements within the CNS can ultimately manifest into distinct disease phenotypes (Larsen et al. 2017).

**Figure 3.**
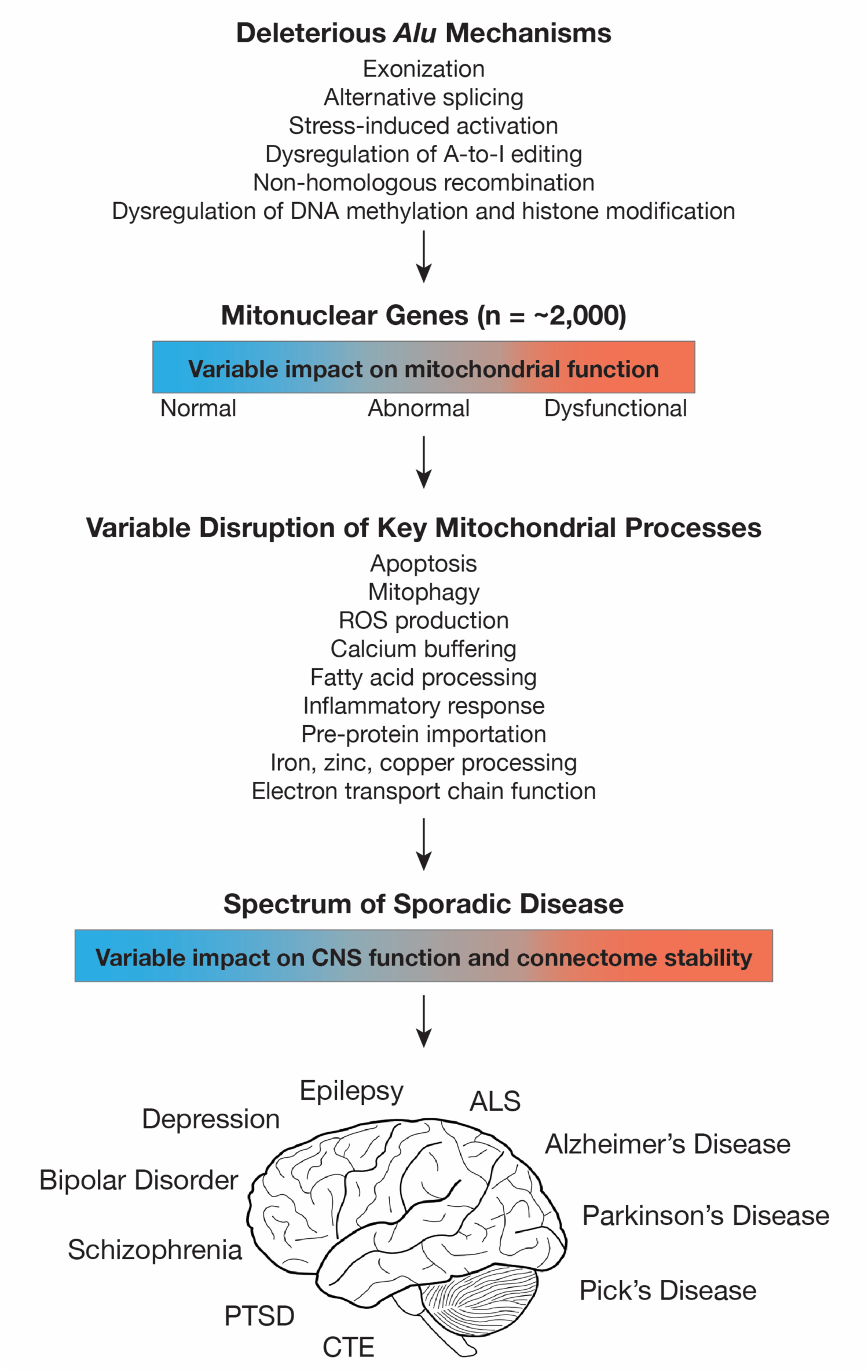
Deleterious *Alu* activity operating on mitonuclear genes can disrupt mitochondrial function in the CNS and contribute to a number of diseased phenotypes (see Table 1). The type and severity of associated neurological and neurodegenerative disorders depends on the deleterious *Alu* mechanism of action, the mitonuclear gene pathways involved, the time or developmental stage of induction, level or severity of traumatic stress, and tissue specificity (see Larsen et al. 2017). If operating across the suite of mitonuclear genes through epigenetic pathways, the mechanism helps to explain the origin of incipient mitochondrial stress and CNS connectome destabilization that is observed across the spectrum of neurological and neurodegenerative disorders.

Several interesting patterns emerge when examining the key neurologic processes that are disrupted through deleterious *Alu* activity (Table 1). For example, mitochondria play an essential role in maintaining intra-cellular metal ion homeostasis (e.g., iron, copper, and zinc), the disruption of which can result in the increased production of free radicals that damage mitochondria and contribute to the increased production of reduced oxygen species (ROS; Rossi et al. 2004; Madsen and Gitlin 2007). The brain is especially sensitive to ROS production, and iron, copper, and zinc-related oxidative stress has been linked to many neurodegenerative disorders including AD, PD, and Wilson’s disease (Rossi et al. 2004; Madsen and Gitlin 2007). It is notable that deleterious *Alu* activity has been identified in several genes that are essential to maintaining proper iron and copper homeostasis, including *FXN*, *ATP7A*, *ATP7B*, *HMBS*, *NDUFS2*, *SLC30A6*, and *PARK7* (*DJ-1*) (Table 1; Gu et al. 2007; Kaler 2011; Girotto et al. 2014). Knowing this, it is possible that either global or tissue-specific dysregulation of *Alu* elements within mitonuclear genes can alter mitochondrial metal ion processing pathways thereby contributing to increased ROS production leading to neurologic stress.

A second interesting pattern with respect to deleterious neurologic *Alu* activity concerns metabolic pathways. The efficient processing of glucose and lipids across the CNS is critical for the stability and function of neurons, and the disruption of mitochondrial-mediated metabolic pathways has been linked to many neurologic disorders including AD and peripheral neuropathies (Viader et al. 2013; De La Monte and Tong 2014). Deleterious *Alu* activity occurs in genes that are critical for glucose and lipid metabolism, including *ABCD1*, *ACAT1*, *ALMS1*, *APOB*, *GK*, *GLA*, *HPRT*, *LPL*, *PDHA1*, *PMM2*, *PSEN1*, *SOD2* and *SPAST* (Table 1). Several of these genes encode for mitochondrial-related proteins that have been implicated in metabolic diseases that directly, or indirectly, contribute to neurological dysfunction. The connection between *Alu* elements and metabolic pathways is consistent with the observation that *Alu* elements preferentially insert into metabolic genes, and this has led to the hypothesis that *Alus* regulate the expression of genes related to Type 1 Diabetes (Grover et al. 2003; Mirza et al. 2014; Kaur and Pociot 2015). Moreover, *Alu* RNAs act to suppress the expression of both endothelial nitric oxide synthase (eNOS) and superoxide dismutase 2 (SOD2) during hyperglycemic conditions (Wang et al. 2016), suggesting a regulatory role of *Alu* elements during oxidative stress and strengthening the link between *Alu* element activity and diabetes.

There is growing evidence linking sporadic AD with dysfunctional metabolic pathways, leading some to consider AD as a ‘Type 3 Diabetes’ wherein glycolysis and lipid homeostasis is altered (Steen et al. 2005; De La Monte et al. 2006; De La Monte and Wands 2008; De La Monte and Tong 2014; De Felice and Lourenco 2015; Mittal et al. 2016). The most well-documented risk factor for AD is a variant within *APOE* (*APOE* ε4), a gene which encodes for a glycoprotein that that mediates cholesterol and lipid transport (Saunders et al. 1993; Strittmatter et al. 1993; Mahley and Rall 2000). The *APOE* ε4 allele is strongly associated with earlier onset of AD, and it is hypothesized that this is a result of the disruption of cholesterol processing and subsequent accumulation of amyloid precursor proteins (APP; i.e., the Amyloid cascade hypothesis). Although the ‘Amyloid cascade hypothesis’ has dominated Alzheimer’s research for decades (Hardy and Higgins 1992; Selkoe 2000; McKhann et al. 2011), the failure of multiple drug trials targeting amyloid pathways has led many in the Alzheimer’s research community to search for alternative hypotheses that can help explain the origin of neurodegenerative disease as well as novel molecular pathways with therapeutic potential (Herrup 2015).

It is of particular interest then to note that a second genetic risk factor for AD, *TOMM40*, is located immediately adjacent to *APOE* on human chromosome 19, and the two genes are in tight linkage disequilibrium (Lyall et al. 2014; Roses et al. 2016a). *TOMM40* encodes for a beta-barrel protein that ultimately forms a central pore in the outer mitochondrial membrane (Shiota et al. 2015) and, much like *APOE*, genetic variants of *TOMM40* are linked to cognitive impairment and neurodegenerative disease (Roses 2010; Gottschalk et al. 2014; Greenbaum et al. 2014; Roses et al. 2016b; Arpawong et al. 2017). The most-well known of these *TOMM40* variants is the rs10524523 (rs523) homopolymer repeat, a variable stretch of deoxythymidine (T) located within *TOMM40* intron 6 (Roses 2010). The rs523 poly-T varies in length from approximately 12 to 46 nucleotides, and the longer variants are statistically associated with thinning of the hippocampus (independent of the *APOE* ε4 allele; Burggren et al. 2017) and earlier onset of AD (Lutz et al. 2010; Roses et al. 2010). Interestingly, rs523 is embedded within tandemly repeated *Alu* elements and originated from an *Alu* insertion event (Larsen et al. 2017). At least 149 *Alu* A-to-I editing events are identified within *TOMM40*, the majority of which are associated with *Alu* elements surrounding the rs523 repeat and intron 9 (Picardi et al. 2017a).

There is a potentially important link that unties *APOE* APP processing with the functional mechanics of pre-protein transport through the TOMM pore. It is possible that conformational changes of the Tom40 protein, potentially originating from *Alu*-mediated events (see above, reviewed in Elbarbary et al. 2016; Chen and Yang 2017; Larsen et al. 2017), can ultimately serve to restrict the passage of lipids across the outer-mitochondrial membrane (Larsen et al. 2017). When combined with altered APP processing, this process could account for the initial site of intra-cellular protein accumulation that is hypothesized to precede extra-cellular plaque formation during very early stages of AD (Skovronsky et al. 1998; D’Andrea et al. 2001; Takahashi et al. 2002). Consistent with this hypothesis is the direct observation of APP accumulation at the TOMM pore (Devi et al. 2006) as well as functional tolerance of Tom40 conformational changes by mitochondria (Mager et al. 2011; Kuszak et al. 2015). Importantly, this mechanism could help to explain the common patterns of protein accumulation (e.g., amyloid plaques and alpha-synuclein Lewy bodies) observed across the spectrum of neurodegenerative disease, including ALS, AD, and PD (Ross and Poirier 2004; Gottschalk et al. 2014; Larsen et al. 2017). An age or stress-related component to Tom40 conformational changes comes with the epigenetic dysregulation of *Alu* elements associated with the aging process or traumatic stress (see Larsen et al. 2017). Whether or not these processes are directly mediated by deleterious *Alu* events remains to be tested, however, it is notable that *Alu* exons and *Alu* somatic retrotransposition events have been identified in several TOM genes that are required for the stability of the translocase of the outer mitochondrial membrane and pre-protein import, including *TOMM5*, *TOMM7*, *TOMM22*, *TOMM40*, and *TOMM40L* (Baillie et al. 2011; de Andrade et al. 2011; Lin et al. 2016).

With respect to *Alu* elements, mitochondrial dysfunction, and the broader pathological scope of AD and other neurodegenerative diseases, there is evidence suggesting that *Alu*-derived peptides interact with tau proteins, perhaps serving a regulatory role for tau phosphorylation (Hoenicka et al. 2002). Tau is a microtubule associated protein that functions to stabilize axonal microtubules and to transport mitochondria along axons, and taupathies (including tau hyperphosphorylation) are a characteristic feature of several neurodegenerative diseases including AD, progressive supranuclear palsy, corticobasal degeneration, and Pick’s disease (Ittner and Götz 2011; Khanna et al. 2016). The *MAPT* gene encodes for tau and alternative splicing events of *MAPT* result in multiple tau isoforms (Reddy 2011). Approximately 86 *Alu* elements (including FLAMs) are distributed throughout *MAPT* introns and A-to-I editing is occurring at 315 *Alu* related sites with elevated levels at the 3’ end of *MAPT* (REDIportal database; Picardi et al. 2017a). When considering the potential for *Alu* structural variants of *MAPT* (including DNA and pre-mRNA secondary structures) and evidence of *Alu* RNAs interacting with tau proteins (Hoenicka et al. 2002), we recommend additional studies aimed at elucidating the regulatory impacts that *Alu* elements might have on *MAPT* gene expression and tau phosphorylation.

### A-to-I editing and the potential for mitochondrial stress

Although several neurological disorders are hypothesized to be the result of disruptive A-to-I editing processes across the CNS (see above), it is presently unknown whether or not these processes are actively influencing mitochondrial function. What evidence is there indicating that post-transcriptional modification of mitonuclear genes can alter gene expression or function? Are there particular neurological or neurodegenerative disorders that are associated with mitonuclear genes that have elevated levels of A-to-I editing? To provide insights into these questions, we searched the REDIportal A-to-I editing database (Picardi et al. 2017a) for mitonuclear genes where 1) A-to-I editing has been identified within *Alu* elements in coding regions and 2) A-to-I editing has contributed to non-synonymous amino acid changes. We identified 57 mitonuclear genes with A-to-I editing occurring within putative *Alu* exons and in 52 of these genes the post-transcriptional modification resulted in nonsynonymous amino acid changes (Supplementary Table 1). Many of these genes are involved with essential neuronal processes including calcium binding and transport, zinc transport, apoptosis regulation, voltage-gated ion channels, and mitochondrial elongation with notable examples including *ADSL*, *BAX*, *CASP2*, *COQ2*, *DFFB*, *FBXO18*, *LYRM4*, *PACRG*, and *SLC30A6* (Supplementary Table 1).

From a broader perspective, we identified enhanced A-to-I editing across 134 mitonuclear genes that are associated with a spectrum of neurologic and neurodegenerative disorders ranging from depression, tobacco use disorder, and bipolar disorder to ALS, Leigh syndrome, PD, and AD (Supplementary Table 2). In light of these patterns, we hypothesize that system-wide or tissue-specific epigenetic dysregulation of *Alu* A-to-I editing within the CNS can serve to disrupt key mitochondrial biochemical processes, thus potentially contributing to incipient mitochondrial and neuronal stress (Figure 3).

## Conclusions

Enhanced somatic retrotransposon throughout neurogenesis contributes to the mosaic brain, however, such activity likely contributes to mosaic pathways leading to disease (Erwin et al. 2014). Elucidating these pathways might ultimately provide insight into the sporadic nature of idiopathic diseases that are impacting the global human population. The disruption of *Alu*-mediated pathways that underlie gene regulation is a plausible mechanism for the origin of complex human-specific neurologic and neurodegenerative disorders. Although many of these disorders have similar phenotypes (e.g., mitochondrial dysfunction), it is possible that these phenotypes arise from deleterious activity operating across tissue-specific gene networks. If correlated with eroding or fluctuating epigenetic control mechanisms of retrotransposons that are associated with aging, cellular senescence, and/or cellular stress (Belancio et al. 2010; Pal and Tyler 2016; Schneider et al. 2017), then such mechanisms might largely escape purifying selection and would be difficult to detect using traditional methods (e.g., genome-wide association studies). It is important to note that the *Alu*-centric mechanisms discussed herein collectively provide a unified framework for multiple hypotheses that have been put forth regarding the origin of neurodegenerative disease including inflammation, oxidative-stress, metabolic dysfunction, and accumulation of protein bodies (see above).

*Alu* elements have played a pivotal role in the evolution of the human epigenome (Prendergast et al. 2014), and both hyper-and hypomethylation of *Alu* elements have been correlated with a number of age-related disorders including Alzheimer’s disease, multiple sclerosis, osteoporosis, and many forms of cancer (Bollati et al. 2009; Jintaridth and Mutirangura 2010; Belancio et al. 2010; Jintaridth et al. 2013; Neven et al. 2016). In light of these patterns, as well as the newly discovered regulatory roles of *Alu* elements (Polak and Domany 2006; Chen and Carmichael 2008; Chen and Yang 2017), we recommend additional research that focuses on the epigenetic interplay between *Alu* elements and mitochondrial gene networks in the central nervous system.

## Acknowledgments

We thank A.D. Brown, W.K. Gottschalk, and M. Mihovilovic for helpful discussion. C. Hohoff, B. Ciapa, and an anonymous reviewer kindly reviewed the manuscript and provided helpful comments. This is Duke Lemur Center publication number 1389.

